# Body Mass Index-Specific Nanoparticle Protein Corona Signatures in Late Pregnancy

**DOI:** 10.64898/2026.05.23.727413

**Authors:** Samantha Velazquez, Matthew C. Juber, Michele Okun, Edward Lau, Ali Akbar Ashkarran

## Abstract

The protein corona (PC) formed on the surface of nanoparticles (NPs) upon exposure to human biofluids is a dynamic interface that reflects the physiological and pathological status of the host. In this study, we investigated how maternal body mass index (BMI) influences the composition of the NPs’ PC during late pregnancy. Polystyrene NPs were incubated with plasma samples collected from third-trimester pregnant individuals across normal weight, overweight, and obese BMI categories. Comprehensive characterization using dynamic light scattering (DLS), zeta potential measurements, and transmission electron microscopy (TEM) confirmed BMI-dependent differences in PC thickness and colloidal stability. SDS-PAGE and label-free quantitative proteomics revealed distinct molecular compositions: PCs from obese individuals were enriched in inflammatory and lipid metabolism-associated proteins (e.g., APOE, CRP), while normal weight-derived PCs showed higher levels of complement regulators and extracellular matrix proteins. Principal component analysis (PCA) demonstrated clear clustering of proteomic profiles by BMI group, suggesting BMI-specific PC fingerprints.

These findings indicate that maternal metabolic phenotype shapes nano–bio interactions at the proteomic level and highlight the potential of PC profiling as a non-invasive approach for assessing maternal health and metabolic status. This work lays the foundation for integrating NP-based proteomics into precision nanomedicine for maternal–fetal health monitoring.

## Introduction

The interface between nanomaterials and biological systems is dominated by the spontaneous formation of a protein/biomolecular corona (PC) —a dynamic layer of adsorbed biomolecules, primarily proteins—upon nanoparticles (NPs) exposure to biological fluids.^1-5^ The composition of the PC layer dictates the biological identity, fate, biodistribution, and cellular interactions of NPs in vivo, ultimately influencing their diagnostic or therapeutic potential.^6-8^ The composition of this complex PC layer is not merely a passive reflection of the plasma proteome; it is influenced by numerous factors including but not limited to disease state, age, sex, and metabolic condition.^2, 9-10^ Among the various parameters that affect the formation and composition of the NPs’ PC, the biological source is particularly important, as it provides the primary reservoir of proteins and/or potential biomarkers that adsorb onto the surface of the NPs.^11-12^ Moreover, the formation of PC is a dynamic and complex process which usually involves competitive adsorption and is influenced by several key parameters including protein concentration, binding affinities, and physicochemical properties of NPs.^1, 13-14^ In this regard, NPs PC has the potential to enrich human plasma proteome due to the dynamic nature of the PC formation on the surface of the NPs.^15^ One of the key aspects of PC in biomarker discovery is enrichment of low abundance biomarkers, which are present in human plasma at very low concentrations and can become enriched in the PC layer.^16-18^ This enrichment facilitates the detection and identification of rare biomarkers that might be challenging to detect directly from the complex biological matrix (i.e., human plasma).^19-20^

Pregnancy represents a unique physiological state characterized by profound immunological, hormonal, and metabolic adaptations that can significantly alter the circulating proteome.^21-23^ Body mass index (BMI), an indicator of nutritional status and metabolic health, further modulates these changes.^24-25^ For example, elevated maternal BMI has been linked to increased risk of gestational diabetes, hypertensive disorders, inflammation, and altered fetal programming.^26-29^ Previous research in the field has explored disease-specific PC patterns, such as those associated with cancer, cardiovascular disease, or neurodegeneration.^30-34^ For instance, NP-assisted proteomic profiling has been used to enrich low-abundance disease-related biomarkers by leveraging the selective adsorption tendencies of NPs.^35^ While NP-assisted proteomic profiling has been successfully used to enrich disease-specific, low-abundance biomarkers via the ‘nano□concentrator’ effect in conditions such as Alzheimer’s disease, breast cancer, and diabetes, there is limited application of this strategy to dissect physiological and metabolic diversity during pregnancy.^36-37^

Although the PC has been extensively studied in disease contexts such as Alzheimer’s and cancer, its responsiveness to physiological variables like BMI during pregnancy remains underexplored.^38^ Our group and others have demonstrated that the PC can serve as a ‘fingerprint’ reflective of the donor’s physiological and pathological state.^32, 39-41^ Building on this foundation, the current study aims to determine whether NPs’ PC composition varies systematically with maternal BMI in the third trimester of pregnancy. By incubating commercially available polystyrene NPs with plasma from pregnant individuals stratified by BMI and subsequently performing label-free quantitative liquid chromatography–mass spectrometry (LC-MS)-based proteomics, we try to uncover differential adsorption patterns, enriched pathways, and potential biomolecular signatures that reflect BMI-related physiological states. This study is designed not only to enhance our understanding of the interplay between metabolic health and NPs-bio interface during pregnancy but also to open new avenues for using PC analysis as a minimally-invasive diagnostic or risk assessment tool. Additionally, the findings may inform NPs design principles for safer and more effective use in pregnant populations, especially in light of the growing interest in nanomedicine applications during pregnancy.

### Experimental details

#### Materials

Pregnant human plasma samples collected from healthy volunteer donors in the third trimester of pregnancy under informed consent and institutional ethical oversight were diluted to a final concentration of 55% using phosphate buffer solution (PBS, 1X). Plain polystyrene NPs (∼ 90 nm) were provided by Polysciences. (www.polysciences.com). Ammonium bicarbonate, urea, dithiothreitol, iodoacetamide, and acetonitrile were purchased from Sigma-Aldrich. Trypsin was obtained from Promega, and formic acid was purchased from Thermo Scientific.

#### PC formation on the surface of the NPs

For PC formation, NPs were incubated with diluted (55%) plasma samples from pregnant individuals so that the NPs’ and human plasma final concentrations were 0.2 mg/ml and 55% in a 1 ml batch, respectively. The solution was then mixed, vortexed well, and incubated for 1h at 37 °C at a constant 1000 rpm agitation. To remove unbound and plasma proteins only loosely attached to the surface of NPs, protein-NPs complexes were then centrifuged at 17,000g for 20 minutes, the collected NPs’ pellets were washed twice more with cold PBS under the same conditions, and the final pellet was collected for preparation for further analysis.

#### LC-MS/MS sample preparation

The NPs’ PC pellets at various BMIs obtained from the various BMI groups were subjected to a standardized proteomic digestion protocol optimized for mass spectrometry based on our previous findings.^42^ Each PC-coated NPs’ pellet was resuspended in 25□µL of buffer containing 50□mM ammonium bicarbonate and 1.6□M urea to facilitate protein solubilization and denaturation. Reduction was achieved by adding 2.5□µL of dithiothreitol (DTT, Pierce BondBreaker, neutral pH) to a final concentration of 10□mM. Samples were incubated at 37□°C for 60 minutes using a Thermomixer. Following incubation, samples were brought to room temperature, vortexed briefly, and centrifuged to collect the contents. Alkylation was performed by adding 2.7□µL of freshly prepared iodoacetamide (IAA) to a final concentration of 50□mM followed by incubation in the dark at room temperature for 1 hour. To quench excess IAA, 3□µL of DTT (10□mM final) was added, followed by a 15-minute incubation at room temperature. Subsequently, sequencing-grade trypsin (Promega), diluted in protein digestion buffer, was added at a 1:50 trypsin-to-protein ratio. Enzymatic digestion proceeded overnight (∼16□h) at 37□°C in a Thermomixer. The next day, the digested samples were cooled to room temperature and centrifuged at 17,000□×□g for 20 minutes to pellet residual NPs. Supernatants containing the tryptic peptides were carefully transferred into fresh low-binding Eppendorf tubes. Formic acid (FA) was added to a final concentration of 5% (v/v) to acidify the samples and adjust pH to between 2 and 3, ensuring peptide stability and compatibility with desalting. Peptide desalting and cleanup were performed using C18 StageTips. Each tip was first activated with 80% acetonitrile (ACN) and 5% FA, equilibrated with 5% FA, and then loaded with the peptide-containing samples. Bound peptides were washed with 5% FA and eluted in two stages: first with 30% ACN + 5% FA, and then with 80% ACN + 5% FA. Both eluates for each sample were pooled, gently mixed, and transferred into a new low-binding tube. Samples were dried using a vacuum centrifuge (SpeedVac) and stored at –20□°C prior to shipment. Dried peptides were shipped overnight on dry ice to the core proteomics facility using FEDEX with next-morning delivery. Sample integrity upon arrival was confirmed by the receiving facility. **Physicochemical characterizations**. DLS and zeta potential analyses were performed to measure the size distribution and surface charge of the nanoparticles before and after PC formation using a Zetasizer Lab series DLS instrument (Malvern company). A Helium Neon laser with a wavelength of 632 nm was used for size distribution measurement at room temperature. TEM was carried out using a JEM-120i (JEOL Ltd.) operated at 120kV. 20 μl of the bare NPs were deposited onto a copper grid and used for imaging. For PC-coated NPs, 20 μl of sample was negatively stained using 20 μl uranyl acetate 1%, washed with DI water, deposited onto a copper grid, and used for imaging. PC composition was also determined using LC-MS/MS.

#### LC-MS/MS analysis and proteomics data processing

Each dried vial was reconstituted with 22 µl 2% ACN 0.1% FA. 5 µL of each digest was injected and run 3 times by nanoLC-MS/MS using a 2h gradient on a Waters CSH 0.075mm x 250mm C18 column feeding into a Thermo Eclipse Tribrid Orbitrap mass spectrometer run in FT-IT mode, using typical settings with HCD for fragmentation. Raw mass spectrometry .raw files were converted to the mzML format using ThermoRawFileParser v.1.4.3. Database searches were performed using Sage v.0.14.7 against the human UniProt Swiss-Prot database downloaded from UniProt in September 2025.^43^ Precursor and fragment ion tolerance were set to ±20□ppm with the following modifications: carbamidomethylation of cysteine (+57.0215□Da, static) and oxidation of methionine (+15.9949□Da, variable). Peptides and proteins were filtered at a 1% FDR threshold. Isoform collapsing and protein parsimony were performed using a custom R-based workflow: peptides mapping to canonical and non-canonical isoforms were collapsed to the canonical protein if not uniquely identifying an isoform, and shared peptides were removed when a non-canonical isoform had a unique peptide. Peptide-level intensities were aggregated to protein-level by summing across peptides, followed by variance stabilization normalization (VSN) using the limma package.^44^ One of the mass spectrometry experiment files contained no protein (NW2 Rep 1) and was removed as an outlier. Principal component analysis (PCA) was performed on normalized protein intensities of the remaining proteins to visualize sample relationships. Samples with low total signal or designated as technical controls were excluded from downstream analysis. Differential expression was conducted using limma v.3.64.3 in R v.4.5.1 with a design matrix with BMI group (normal weight, overweight, and obese) and subject sample as factors.^45^ Significance was determined using a limma FDR-adjusted p-value cutoff ≤ 0.05 and absolute log_2_FC ≥ 1. Additional analysis was performed with the aid of the ggplot, clusterProfiler, and pheatmap packages in R.

## Results and discussion

An overview of the experimental workflow is illustrated in **Figure 1**. Plasma from third-trimester pregnant women across BMI categories (normal weight, overweight, and obese) was incubated with polystyrene NPs, followed by isolation/purification, characterization, and proteomic analysis to uncover compositional differences in the resulting PC composition. In this study, commercially sourced plain polystyrene NPs with a mean diameter of ∼ 90□nm were used as the model nanomaterial. Polystyrene NPs are extensively used as model systems in nanomedicine and clinical research due to their well-established biocompatibility, chemical stability, and low toxicity profile.^7, 46^ Therefore, PSNPs offer a robust platform to understand how host factors, such as maternal BMI, modulate the bio–nano interfaces which is an essential insight for developing personalized nanomedicine strategies.^47-48^

**Figure 1:**
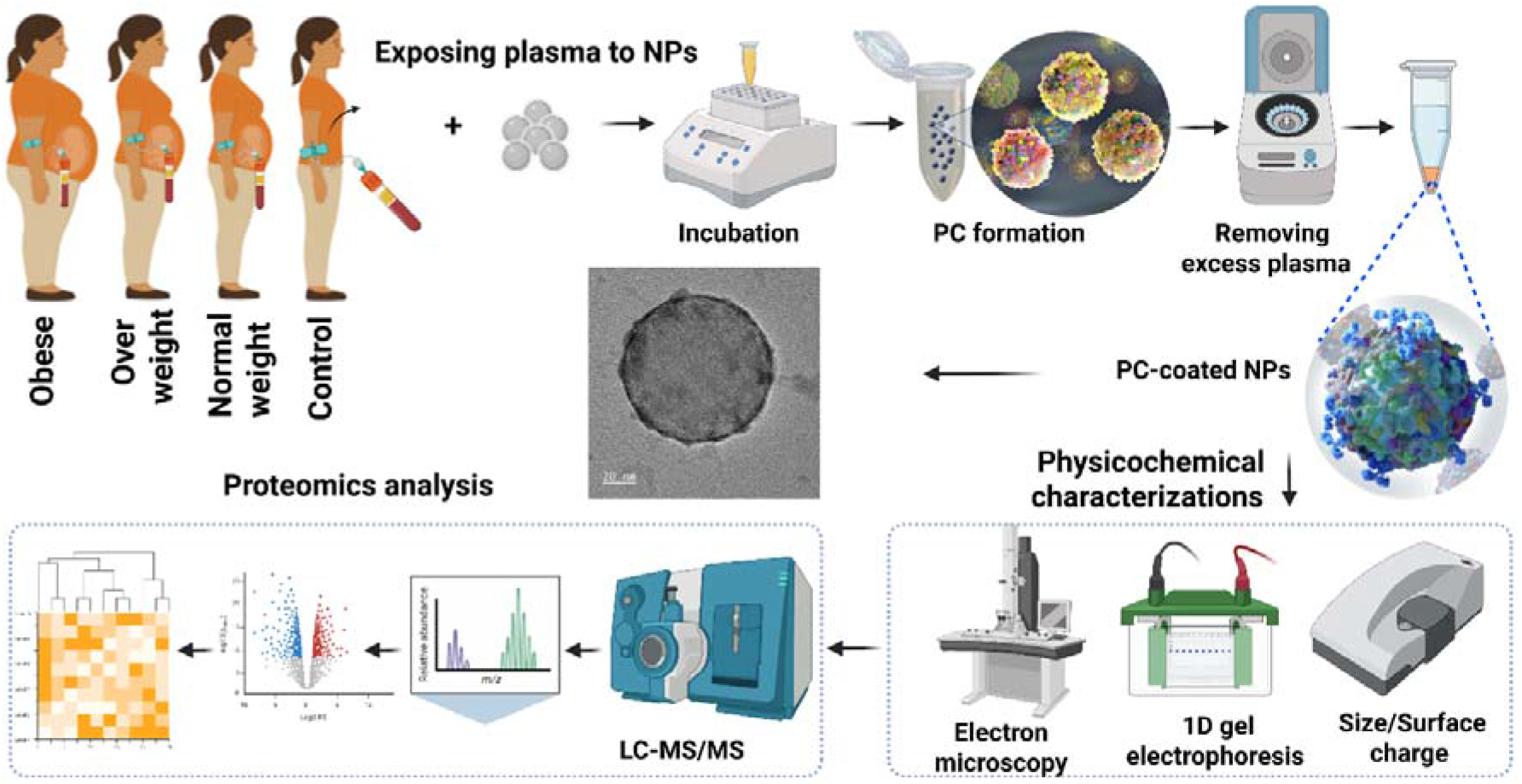
Schematic overview of the experimental workflow for identifying BMI-specific NPs’ PC signatures in late pregnancy.

The physicochemical characterization of both uncoated (bare) and PC-coated NPs was conducted using DLS, zeta potential measurements, and TEM. The bare polystyrene NPs were monodispersed with a narrow size distribution, exhibiting an average hydrodynamic diameter of ∼ 93□nm and a surface zeta potential of ∼ –35□mV (**Figure 2** and **Table 1**). Upon incubation with pregnant human plasmas, the NPs underwent noticeable surface remodeling: the average size increased to about 100-130 nm and the surface charge shifted to –15 to -10□mV approximately, reflecting the successful formation of a PC layer. Three different BMI groups are denoted by NW for normal weight, OW for overweight, and OB for obese and numbers 1 to 3 refers to three different individual samples within each BMI group.

**Table 1:** Average size, polydispersity index (PDI), zeta potential, and the corresponding standard deviation values of bare NPs and PC-coated NPs at various BMIs (three repeats).

| NPs | Size (nm) | SD (nm) | PDI | SD | Zeta potential (mV) | SD (mV) |
| --- | --- | --- | --- | --- | --- | --- |
| **Bare** | 92.8 | 0 | 0.055 | 0.005 | -35.0 | 0.6 |
| **Control** | 132.4 | 9.6 | 0.354 | 0.015 | -12.3 | 0.6 |
| **NW 1** | 133.4 | 17.9 | 0.321 | 0.007 | -15.7 | 0.1 |
| **NW 2** | 119.7 | 8.3 | 0.204 | 0.031 | -14.6 | 0.3 |
| **NW 3** | 122.6 | 4.1 | 0.258 | 0.021 | -10.5 | 1.2 |
| **OW 1** | 110.9 | 4.1 | 0.150 | 0.014 | -13.3 | 0.7 |
| **OW 2** | 100.4 | 6.1 | 0.324 | 0.004 | -15.5 | 0.9 |
| **OW 3** | 113.8 | 8.3 | 0.208 | 0.007 | -10.2 | 0.9 |
| **OB 1** | 113.8 | 8.3 | 0.245 | 0.051 | -12.3 | 0.4 |
| **OB 2** | 105.4 | 3.5 | 0.164 | 0.005 | -11.4 | 0.3 |
| **OB 3** | 119.7 | 8.3 | 0.216 | 0.016 | -10.2 | 0.2 |

**Figure 2:**
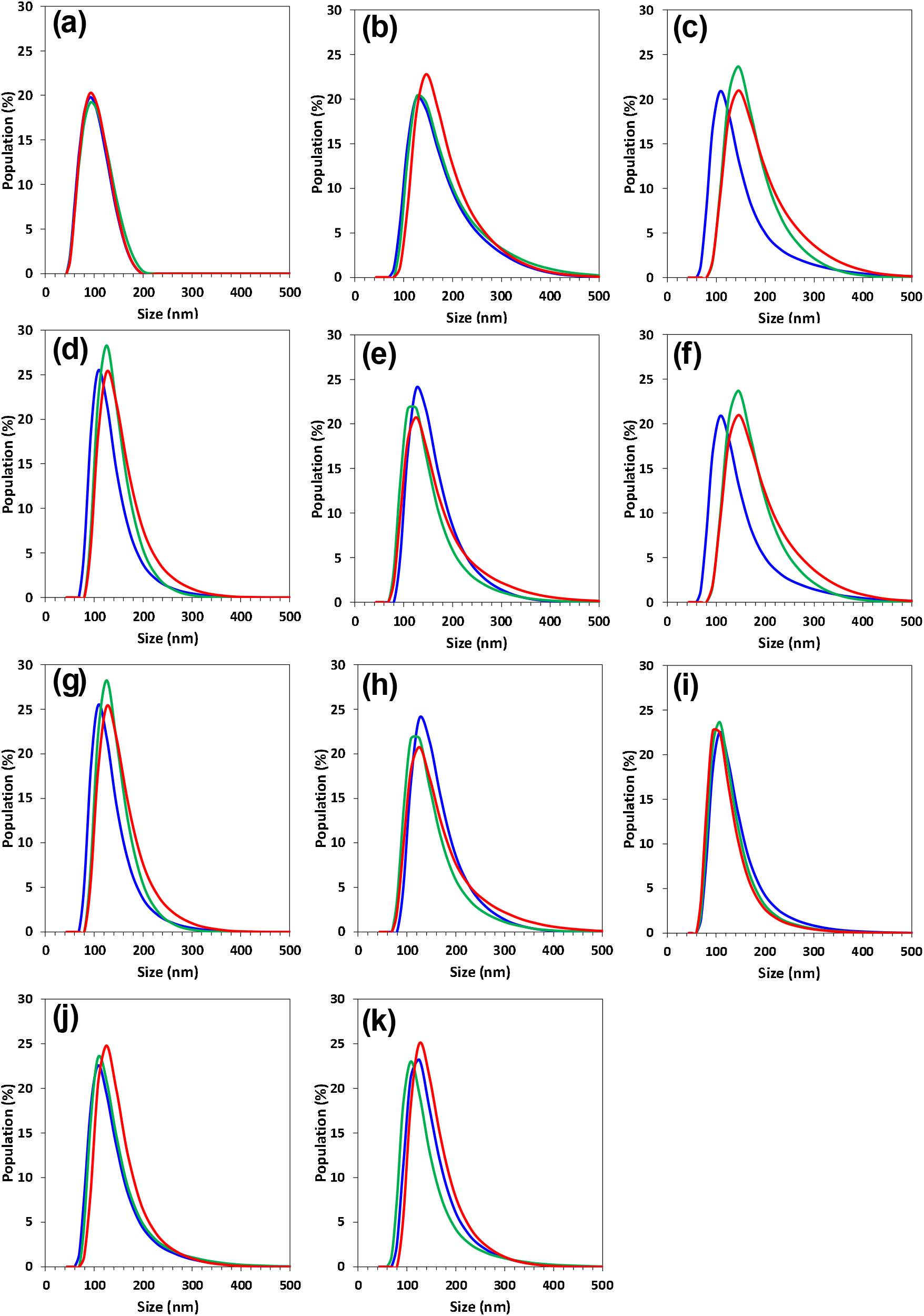
Size distribution of (a) bare NPs, PC-coated NPs of (b) control, (c) NW1, (d) NW2, (e) NW3, (f) OW1, (g) OW2, (h), OW3, (i) OB1, (j) OB2, and (k) OB3 (three repeats).

The average size, polydispersity index (PDI), and surface charge of all samples are summarized in **Table 1**. As shown in **Table 1**, the average sizes and surface charges increased and decreased respectively in all samples upon formation of the PC on the surface of the NPs. TEM imaging provided visual confirmation of these changes, highlighting distinct morphological differences between bare and PC-coated NPs (**Figure 3**). Specifically, a dark halo surrounding the NPs’ core was evident after exposure to human plasmas, indicating the presence of an adsorbed protein layer (**Figure 3c** and **3d**). Microscopic images of the PC coated NPs confirm that a thin dark layer surrounded NPs which indicates that NPs are randomly coated with PC.^49-51^ As is clear from the images bare NPs are highly monodispersed with a narrow size distribution while PC coated NPs revealed presence of a thin layer of proteins on the surface of the NPs. The increased average size and PDI of PC-coated NPs incubated with pregnant plasma samples, as observed in our DLS measurements, are consistent with the morphological changes confirmed by electron microscopy. As there was not a significant difference in TEM images of PC-coated NPs obtained from various BMIs, the sample prepared with NW PC-coated NPs is presented as representative in **Figure 3** (see **Figure S1** of Supplementary Information (SI) for TEM images of PC-coated NPs at various BMIs conditions).

**Figure 3:**
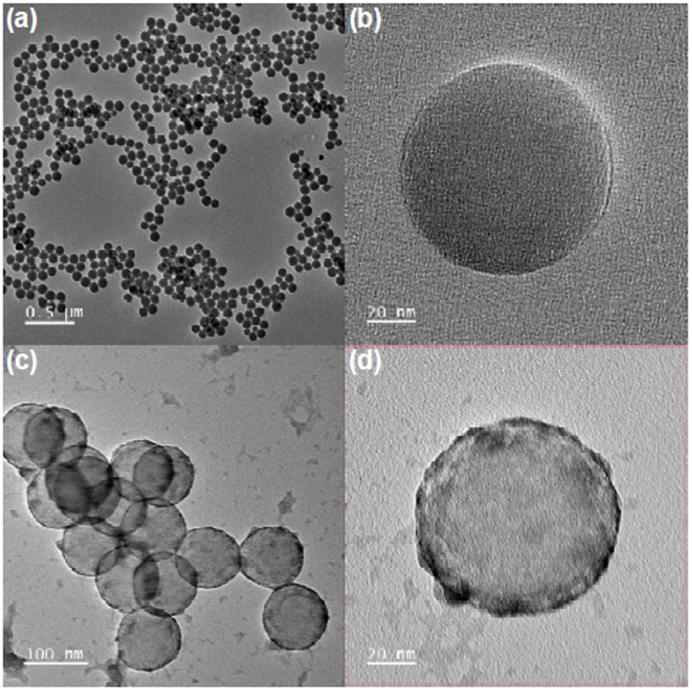
(a, b) TEM images of bare NPs and (c, d) representative TEM images of PC-coated NPs following incubation with NW third-trimester pregnant plasma.

As a next step, we analyzed the PC profile at the surface of the NPs using one-dimensional (1D) SDS-PAGE analysis (**Figure 4**). Although the resolution of the gel electrophoresis analysis is not comparable with mass spectrometry, the findings show distinct banding patterns across various BMIs and control samples. All lanes corresponding to control NPs’-PC and PC-coated NPs prepared at various BMIs exhibited multiple bands across a wide molecular weight range, confirming successful protein adsorption to the NPs’ surface. Between-group differences were observed in both the intensity and distribution of bands, suggesting compositional variability in the PC profile related to maternal BMI. For example, lower molecular weight bands (∼15–25 kDa) appeared more intense in NW samples, whereas higher molecular weight bands were slightly enriched in OB samples, possibly reflecting altered proteomic signatures associated with metabolic state. Moreover, across all BMI categories, SDS-PAGE revealed a conserved core protein profile with subtle differences in band intensity in the ∼25–75 kDa range, possibly indicating enrichment of acute-phase proteins or metabolic enzymes. These qualitative differences were further evaluated through LC-MS/MS proteomic analysis to characterize individual proteins and identify potential BMI-specific PC signatures.

**Figure 4:**
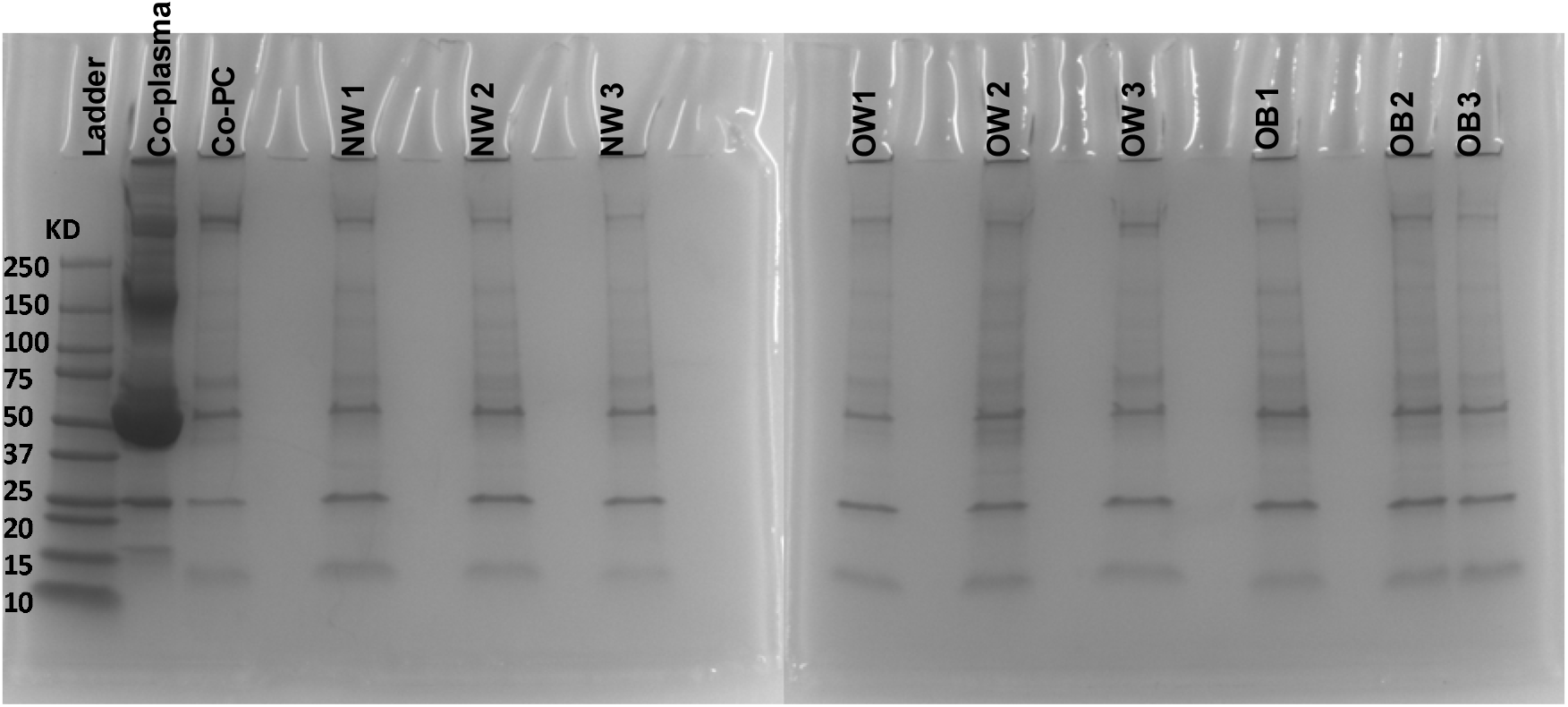
SDS-PAGE analysis of PC compositions from NPs incubated with third-trimester pregnant plasma samples across different BMI groups.

We performed a comprehensive proteomics analysis of the PC profiles of each group, quantifying a total of 644 proteins despite the lack of further fractionation, consistent of an enrichment effect of low-abundance proteins. To quantitatively compare the PC proteome profiles, the mass spectrometry intensities of proteins were normalized using variance stabilization normalization (VSN) with the limma R package ^45^ to improve comparability across various LC-MS runs. Principal component analysis (PCA) of the variance-stabilized protein samples clustered by individual participant samples as presented in **Figure 5a**. PCA analysis clearly demonstrates separation trends between the BMI groups, with obese and normal weight forming more distinct clusters, while overweight partially overlaps with both. This indicates differential PC composition that is influenced by maternal BMI. The protein intensities had comparable overall distributions (**Figure 5b**), highlighting the consistency of the PC profile across participant samples. The clustering patterns reinforce the PCA findings and reveal group-specific protein expression signatures. Several clusters of proteins appear to be enriched or depleted in association with BMI status, indicating potential biomarker candidates. Moreover, the identified protein distribution spanned 5 orders of magnitude, demonstrating the NP’s efficacy in capturing proteins across the dynamic range of the proteome (**Figure 5c**). Additionally, unsupervised hierarchical clustering highlighted a number of distinct proteomic signatures across BMI groups (**Figure 5d**). Samples from overweight and obese participants tended to cluster together, while normal weight samples formed a separate group. This clustering reinforces the idea that even in the absence of clinical pathology, BMI alone can shape the nano–bio interfaces. Proteins contributing to the clustering pattern included vitronectin, complement factor B, and fibronectin, all of which are associated with immune modulation and extracellular matrix remodeling, processes that may be subtly altered across BMI groups.^52^

**Figure 5:**
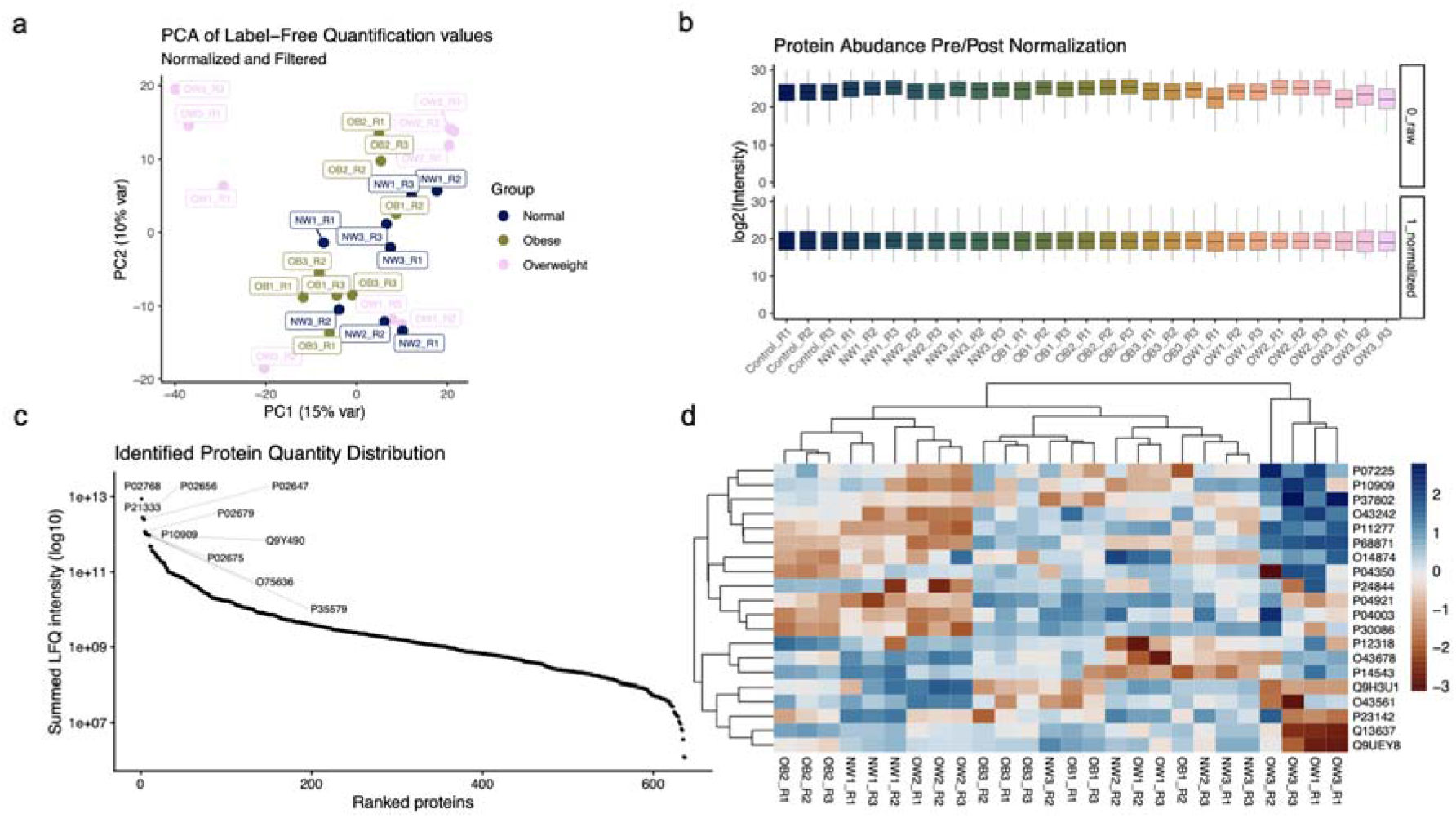
Proteomic profiling of proteins enriched from NPs incubated with third-trimester pregnant plasma samples across different BMI groups. (a) PCA of the intensities of 644 PC proteins from mass spectrometry experiments. (b) Box plots showing protein abundance distribution in each sample before (top) and after (bottom) normalization. (c) Rank plot showing the summed label-free quantitation (LFQ) intensity of proteins measured in the experiment. (d) Heatmap showing top 20 most variable proteins.

Differential expression analysis revealed significant differences in multiple protein families in the PC profiles across BMI categories (**Figure 6a-c**). Several proteins were significantly increased in the obese group relative to normal weight, suggesting a distinct alteration in the plasma protein composition that affects NPs’ surface adsorption (limma FDR adjusted P < 0.05). Notably, Apolipoprotein C3 (APOC3) and serum amyloid A1 (SAA1) were upregulated in the obese group, both of which are markers of dyslipidemia and systemic inflammation, consistent with the known metabolic dysregulation associated with higher BMI during pregnancy.^53-55^ These statistically significant differences strengthen the argument that BMI modulates the PC in a measurable and biologically meaningful way. Additionally, Gene Ontology Biological Process (GO:BP) pathway analysis revealed a significant enrichment (FDR adjusted P < 0.01) of lipoproteins, complement proteins, and coagulation related terms across BMI categories (**Figure 6d-f**). Our findings suggest that apolipoprotein (a), a known atherogenic protein, was markedly elevated in individuals with higher BMI, showing significant increases in both the overweight and obese groups compared with those of normal BMI (log_2_FC = 3.19 and 3.57, respectively).^56-57^ In contrast, apolipoprotein B, a structural component of low-density lipoprotein (LDL) and a strong biomarker of atherosclerotic risk, was specifically elevated in the obese group relative to overweight individuals.^58^

**Figure 6:**
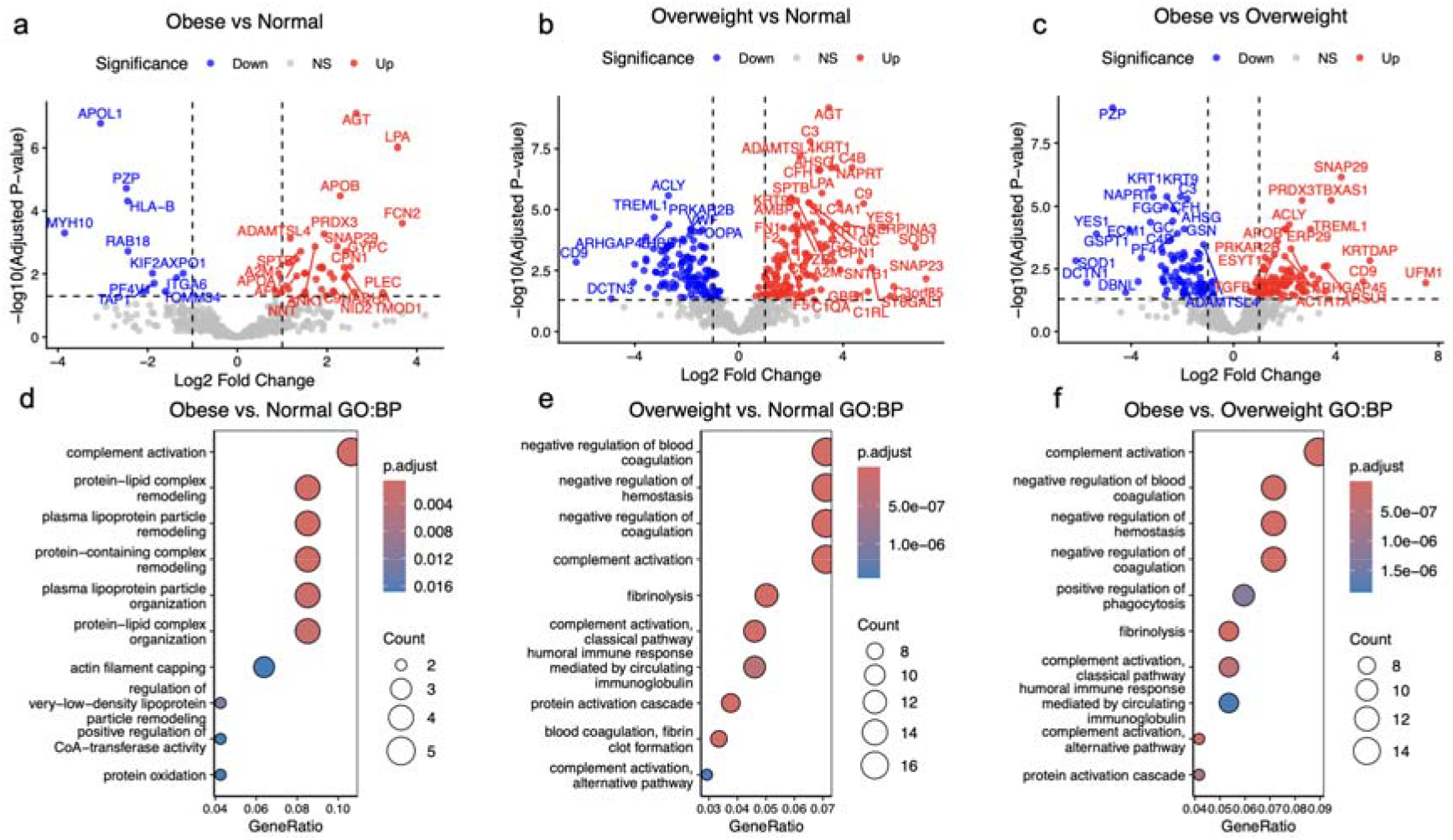
Differential expression analysis of normalized protein abundance compared across BMI groups: (a) Obese vs Normal BMI. (b) Overweight vs. Normal BMI. (c) Obese vs. Overweight BMI. Labeled points met a threshold deemed statistically significant defined by an adjusted p-value < 0.05 and absolute log_2_FC > 1. Gene enrichment for biological processes of statistically significant proteins across comparisons: (d) Obese vs. Normal BMI, (e) Overweight vs. Normal BMI, and (f) Obese vs Overweight BMI.

Notably, carboxypeptidase N catalytic chain (Kininase-1), an enzyme involved with the generation of kinin B1 receptor (BIR) agonists was shown to be increased in overweight and obese groups (log_2_FC = 3.53 and 2.52 respectively). B1R plays an important role in mediating inflammatory response with upregulation having been linked to neuroinflammation and hypertension.^59-60^ Additionally, B1R activation has been shown to contribute to insulin resistance through activation and upregulation of pro-inflammatory cytokines and inducible form of nitric oxide synthase.^61^ These findings align with prior studies showing that maternal obesity promotes low-grade inflammation and alters vascular and metabolic regulation — and for the first time, our data demonstrate that these systemic effects are mirrored at the nano–bio interface level.^62-63^

In parallel, pregnancy zone protein (PZP), an immunosuppressant secreted by the liver in increased levels during pregnancy, was sensitive to changes in BMI.^64^ Notably, PZP interacts with several receptors and chaperone proteins, including lipoprotein receptor-associated protein (LPR), transforming growth factor-β (TGF-β), glycoside A (GdA), and 78kDa glucose-regulator protein (GRP78).^64^ Conflicting reports demonstrate PZP levels as increasing or decreasing during diabetes mellitus, obesity, and some cancers.^64^ Here we find PZP abundance declined progressively with increasing BMI, being lower in obese compared with both normal and overweight individuals (log_2_FC = ™2.47 and ™4.72, respectively), but was modestly higher in the overweight group compared with normal (log_2_FC = 2.25). Additionally, human leukocyte antigen (HLA) antigen B, a key protein involved in self-recognition in the adaptive immune system, demonstrated decreased levels in both overweight and obese individuals (log_2_FC = -1.84 and -2.45 respectively).^65^ Taken together, the proteomics results revealed significant differences in PC proteome profiles across each group, highlighting a number of biomarker candidates for understanding the effect of BMI on pregnancy. The results also reinforce the utility of PC toward the characterization of complex clinically relevant biofluid samples. The full protein list can be found in **Supplementary Table S1**.

## Conclusions

In summary, our study demonstrates that maternal BMI during late pregnancy significantly alters the composition of the NPs’-PC, with reproducible differences in both the abundance and identity of adsorbed plasma proteins across normal weight, overweight, and obese groups.

Through integrated physicochemical characterization and LFQ proteomics, we uncovered distinct molecular fingerprints associated with maternal metabolic phenotype. PCs derived from obese individuals were enriched in proteins involved in lipid metabolism (e.g., APOE, LPA), pro-inflammatory pathways (e.g., CRP, SAA1), and immune regulation—molecular signatures that are consistent with the systemic inflammation and metabolic stress reported in maternal obesity. In contrast, PCs from normal weight pregnancies showed enrichment in proteins related to complement cascade regulation, extracellular matrix remodeling, and homeostatic immune signaling, reflecting a more balanced physiological state. These findings highlight that the nano–bio interface is highly sensitive to metabolic variations in the host and that NPs serve as effective molecular concentrators, capturing clinically relevant proteomic differences. Notably, the observed clustering in PCA and the qualitative shifts in PC composition suggest a strong association between host BMI and PC fingerprinting, supporting the concept of PC-based diagnostics. By capturing BMI-dependent molecular patterns in late pregnancy, our platform paves the way for the development of personalized nanomedicine tools for prenatal risk stratification and biomarker discovery. Future studies leveraging longitudinal cohorts and higher-resolution proteomic techniques are warranted to validate these findings and to explore their relevance in pregnancy-associated disorders such as gestational diabetes, preeclampsia, or fetal growth restriction.

## Associated content

## Supporting Information

Supporting figures/tables, TEM images, and LC-MS/MS analyses.

**Notes**

The authors declare no competing financial interest.

## Data Availability

The raw mass spectrometry data associated with this project have been uploaded to ProteomeXchange via JPOST at PXD070237 (Reviewer access https://repository.jpostdb.org/preview/14403436906909041e1de9b Access key 6771)

## Acknowledgments

S.V and A.A.A acknowledge financial support from National Institutes of Health (grant number R03EB034817). E.L. is supported in part by the University of Colorado School of Medicine Translational Research Scholar Program (TRSP) and the Research Investment in the Scientific Enterprise (RISE) awards.

## Notes

### Competing Interest Statement

The authors have declared no competing interest.

